# Dosimetric Study of Breast Scar Marker in Different Radiotherapy Dose Calculation Algorithms

**DOI:** 10.1101/535476

**Authors:** S. S Abdullah, Ahmed Alsadig, A. Sulieman, Isam H. Mattar, C. K. Ying, K. Gokula, A. N. Azahari, M. Z Abdul Aziz

**Affiliations:** Oncological and Radiological Science Cluster, Advance Medical and Dental Institute, Universiti Sains Malaysia, 13200 Kepala Batas, Penang, Malaysia; Elettra-Sincrotrone Trieste S.C.p.A., 34149, Basovizza, Trieste, Italy; Universita di Trieste, Piazzale Europa, 1, 34127 Trieste, Italy; Radiology and Medical Imaging Department, Collage of Applied Medical Sciences, Prince Sattam bin Abdulaziz University, Alkharj, Saudi Arabia; Department of Radiological Sciences, College of Applied Medical Sciences, King Saud University, P.O Box 10219 Riyadh 11433, Saudi Arabia

**Keywords:** CC and PB algorithm, 3D CRT Breast Cancer, Photon Beam Perturbation

## Abstract

The purpose to this work is to validate and benchmark the delivered dose accuracy during radiotherapy treatment (without marker at scar site) to the dose calculated by treatment planning that included a marker on scar site. Dose distributions in breast cancer 3-dimensional conformal treatment planning (3D CRT) calculated with Pencil Beam (PB) and Collapsed Cone (CC) algorithms of commercial treatment planning system (Monaco and Oncentra MasterPlan) was compared as photon beam through homogenous and heterogenous media (by placing marker on phantom surface) to evaluate the perturbation of photon beam. Radiochromic film dose distribution value was compared to the dose calculated by CC algorithm of Monaco and Oncentra MasterPlan (OMP) Treatment Planning Systems (TPS) and PB algorithm of OMP TPS. During Computed Tomography (CT) simulation procedure for breast case, a metal-based scar marker (wire) was used to localize the tumour bed during treatment planning procedure which the phantom was considered as heterogeneous medium. In homogenous medium, PB algorithm gave smaller dose deviation compared to CC algorithm. When wire was introduced to the surface of phantom, PB algorithm (6.0 cGy) gave higher dose deviation compared to CC algorithm (2.0 cGy). However, CC algorithm (plasticine: 7.0 cGy and cable: 7.3 cGy) shows higher dose deviation compared to PB algorithm (plasticine: 6.3 cGy and cable: 6.6 cGy) when Plasticine and Cable marker were introduced. The placement of marker in lateral orientation gave smaller perturbation to megavoltage photon beam compared to axial orientation in overall case. Moreover, wire and plasticine are suitable as a scar marker due to its tissue equivalent density with less perturbation to photon beam.

## 1. Introduction

Breast malignancy is the most common cancer in females globally and contribute 25% of all cancer incidence. In females, A breast cancer treatment or radiotherapy is started with surgery (mastectomy or lumpectomy) as neoadjuvant treatment, and the remaining microscopic tumour cell (tumour bed) is treated via radiation therapy as an adjuvant treatment for a complete tumour cell death (tumour control) with minimal tissue reactions (normal tissue complication probability (NTCP) with accuracy 5% or better (ICRU 1976). The site of tumor bed is marked at post-surgery scar during computed tomography (CT) simulation. The scar marking is useful in treatment planning three-dimensional conformal radiotherapy treatment (3D CRT) as it displays the extension of remaining microscopic tumour that should be included around beam field.

The characteristics of the marker includes; the appearance in the imaging system used for radiation therapy treatment planning (RTP) such as CT, artifacts produced in an image is minimized, and perturbation of the treatment radiation dose to the clinical target volume (CTV) is minimum [1]. All the above characteristics of a marker can be achieved by a radiolucent marker with a low atomic number and mass density that equivalent to the density of water (1.0 g.cm^-3^). Metal-based marker (as example wire) was commonly used during simulation procedure. This is because the marker has a mass density close to water density despite having high atomic number. However, the calculation of dose between treatment planning and dose delivered during treatment just below the marking site will become significantly different due to perturbation of dose through the marker during dose calculation. The scar marker in CT images data commonly show streaks artifact and cause the dose distribution underestimate or overestimate from the standard dose distribution throughout the depth from surface. Underestimating dose to target will cause insufficient dose to tumor (recurrent cancer cell development) while overestimating dose will insecure nearby organs at risk (in this case; patient’s cardiac muscle).

The dose calculation is performed by collapsed cone algorithm from Monaco TPS and collapsed cone and pencil beam algorithms from Oncentra MasterPlan TPS for 3D CRT. An EBT3 GaFChromic™ film dosimetry was used for a verification of TPS dose calculation. The dose distributions calculated by both algorithms were compared to the measured dose distribution. In this study, the dose distributions between collapsed cone algorithm and pencil beam algorithm was compared in the homogenous (no marker placement) and heterogenous (by placing marker on phantom surface) media. A homogenous phantom was used to avoid other influencing factors such as, heterogeneities effects and contour effects affect the results of the study.

Collapsed cone algorithm is typically used for photon beam as an option to pencil beam dose calculation. The principle of calculation dose using collapsed cone algorithm involves using an input data from a beam model. The beam model is a display of the characteristics of radiation delivery for a machine and energy combination. The beam modelling can be defined as characteristics of the radiation irradiated by an apparatus and combination of nominal energy. The features involved the energy spectrum, output factors, profile definition, and geometric machine characteristics. The next step is the calculation of primary 2-dimensional (2D) energy fluence from the linear accelerator for one field in a period. The calculated fluence energy is then projected pass the patient model. The primary fluence is the amount of photon energy delivered per unit area directly from the linear accelerator machine. After that, a 3-dimensional (3D) distribution of Total Energy Release in Matter (TERMA) is calculated. TERMA is known as the amount of energy released in the patient in a specific area. The final step is the combination of TERMA to a dose spread function for 3D dose deposition calculation [3].

The other dose calculation technique is the pencil beam algorithm. For this type of beam, any collimated incident photon beam that irradiated on a patient is in the form of a group of smaller, narrow “pencil beams.” A central ray axis for each of these pencil beams deposited an amount of dose. The dose deposition modelling differs in term of intensity and spectrum of the beam irradiated on a patient. Fields shapers such as linear accelerator jaws, blocks and multi-leaf collimators are used to define the arrangement and weighting of the pencil beams. The non-uniform or modulated beam profile is produced by adjusting the weighting of each pencil beam. The weighting of each beam involved the primary photon intensity of the entrance dose as well as electron contamination. Then, the total incident photon energy for pencil beam is called the primary energy fluence. In pencil beam, one of the pencil beams is separately considered where it has a minute diameter on the surface. The dose is deposited under the surface when the pencil beams hit the surface. Thus, by following the primary scattering and absorption process, the dose deposited (photons and secondary electrons) will have a definitive spatial distribution and spread out to form a tear-drop or pear-shaped distribution of dose. It is referred to as pencil beam dose kernel or dose kernel [4].

The differences between collapsed cone and pencil beams algorithms are classified in different type. Type A is a type of algorithm where the electron (energy) transport not modelled while type B is a type of algorithm where the electron transport is considered. Pencil beam is a type A algorithm under pencil kernels while collapsed cone algorithm is a type B algorithm under point kernels. In heterogeneous medium, type A algorithm proved be inadequate in heterogeneous medium dose calculation especially for small fields. However, there are no algorithm that able to adequately calculate the dose under all conditions [5].

The purpose to this work are to validate and benchmark the delivered dose accuracy during treatment (without marker at scar site) to the dose calculated by treatment planning that included a marker on scar site. To assess the accuracy of dose for a total fraction of a breast radiation therapy to obtain the total uncertainties of dose delivery. And finally to evaluate the effect of marker placement (marker was placed in axial and lateral orientation) to the dose distribution just below the marker was also investigated.

## 2. Materials and Methods

The study had been conducted using homogenous phantom to study the effect of dose distribution solely on scar marker. The accuracy of dose distribution was evaluated using percentage dose deviation. **Error! Reference source not found**. summarizes the methodology followed in this study.

### 2.1 Phantom Preparation

14 slabs (1 cm each) and 2 slabs (0.5 cm each) of solid water equivalent phantom were arranged to form a 15 cm homogenous phantom as shown in Figure 2(a). Another slab of the phantom with 30 cm height and 30 cm width with its centre marked on phantom as isocentre is shown in Figure 2(b). The combination of all slabs formed a homogeneous phantom. Homogenous phantom was used in this study instead of thorax or chest phantom to avoid other factors such as lung cavity (low density) factor contributed in dose calculation.

### 2.2 Scar Marker Preparation

Three type of scar marker were used in this study, which is a wire (metal-based marker), cable and plasticine as shown in **Error! Reference source not found.**. The density determination kit was used in determining the mass density of each marker. Distilled water with known mass density (1.000 g/cm^3^) and density of air (0.001225 g/cm^3^) were used in weighting procedure. The markers were weighted in the air and inside the distilled water. The mass values obtained were used in mass density calculation:

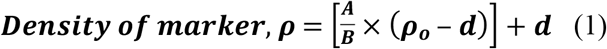

where, A is weight of marker in air (g), B is weight of marker in water (g), d is the density of air and ρ_o_ is density of distilled water. The calculated mass density was used in this study as in **Error! Reference source not found.**.

### 2.3 CT Simulation Scanning

The set-up for CT simulation process is seen in Figure 1. Four sets of CT data were taken using Toshiba Aquilion™ CT Simulator labeled as “control” which has no marker placement on phantom surface, “wire” with wire scar marker placement, “cable” with cable marker placement and “plasticine” with plasticine marker placement. An example of marker positioning is illustrated in **Error! Reference source not found.**.

**Figure 1:**
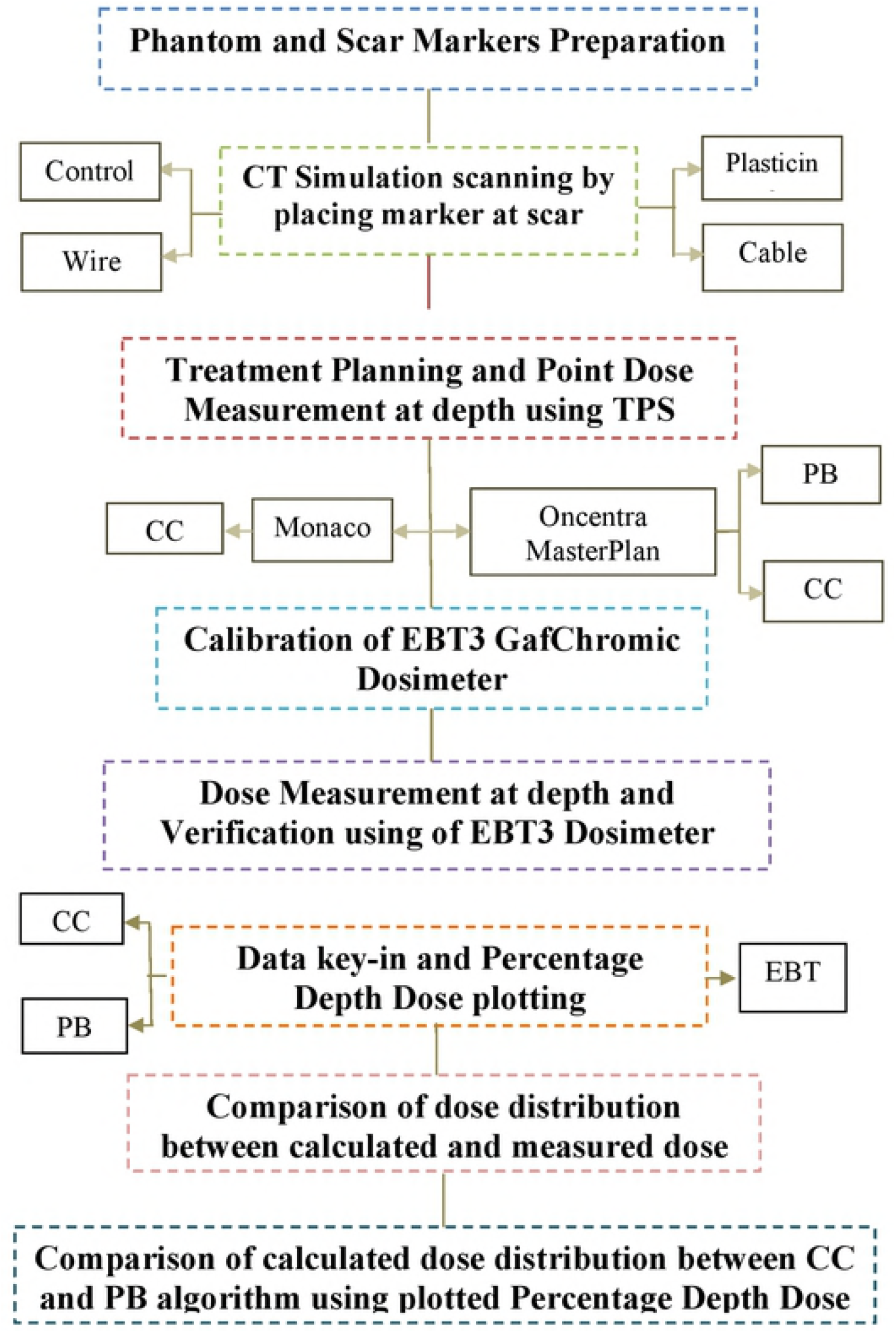
CT simulation set-up of solid water phantom (a) Lateral and (b) superior views.

### 2.4 Treatment Planning and Dose Measurement Using a Treatment Planning System (TPS)

The planning strategy involved two commercial treatment planning systems (TPS), Monaco^®^5 that provided Collapsed Cone (CC) algorithm and Oncentra MasterPlan that provided CC and Pencil Beam (PB) algorithms for 3D CRT. A direct beam of 6 MV photon was used with 15 cm x 15 cm field size to ensure that marker placement within the irradiated area and to avoid significant variation in dose distribution where smaller field area irradiation will show a substantial variation in dose distribution [6, 7]. The standard total prescription dose for breast cancer case used in Radiotherapy Unit is 4256 cGy for 16 fractions. Therefore, the prescribed dose for a fraction is 266 cGy at prescription point (Point A) that used in this study. Point doses were created from the surface to 10 cm depth below the markers (at central axis for “control” CT images) for axial and lateral placement of marker as in Figure 1. The same planning was used for four sets of CT images. Then, three sets of plans with marker placement were modified. The mass density of marker’s contour was corrected with the mass density of air and water. All the dose distribution data were collected and tabulated. The dose at depth curve was plotted.

### 2.5 Calibration of Film Dosimeter

An EBT3 radiochromic film was cut into 11 pieces with dimensions of 3 cm × 3 cm and labeled. The films were placed at Point A (1.5 cm depth) of phantom as shown in **Error! Reference source not found.**. The dose used to irradiate the films as in **Error! Reference source not found.**. All films were scanned using EPSON Expression 11000xl Flatbed Scanner and read using FilmCal software. The calibration curve is shown in **Error! Reference source not found.**.

### 2.6 Dose Measurement and Verification Using Film Dosimeter

Another EBT3 film was cut at 30 cm× 32 cm and placed between phantom slabs as in **Error! Reference source not found.**. 15 x 15 cm^2^ field size was used with 253 MU.

### 2.7 Film Scanning and Reading

The films were scanned using EPSON Expression 11000 xl Flatbed Scanner and Film Scan software. The image was read by PTW VeriSoft software. A plotted beam profile was displayed, and normalization point was set at 3422.480 cGy where the depth at 1.5 cm. The tabulated data and plotted PDD curve in absolute and relative data were recorded and saved.

## 3. Results and Discussion

### 3.1 Verification of calculated dose distribution

The results of dose verification are shown by comparing the dose distribution between the dose measured by EBT3 radiochromic film and the dose calculated by algorithms of TPS. **Error! Reference source not found**. and **Error! Reference source not found**. were plotted based on **Error! Reference source not found.**. Based on **Error! Reference source not found**., percentage dose deviation at the surface between the dose measured by EBT3 and dose calculated by CC algorithms of Monaco TPS was 1.4%. A significant deviation is shown by dose distribution calculated using CC algorithm of Monaco TPS from the surface to Point A (prescription point or 1.5 cm from surface) ranging from 0.0% cGy to 1.4%.

The tolerances dose accuracy is 2% or 2 mm in simple cases. In this study, the plan used was direct beam which under simple planning [8]. Therefore, the dose accuracy should be within the tolerance without considering other complexity. Based on the results, the percentage deviations shown by the algorithms were below 2% and within the tolerance limit. Therefore, the calculation of the algorithms was acceptable in this study. The dose distribution at the prescription point calculated by the algorithms has shown no deviation when compared to measured dose distribution. Furthermore, the percentage deviation of dose distribution calculated by the algorithms almost the same as the depth beyond Point A.

### 3.2 Dose distribution between Collapsed Cone and Pencil Beams algorithms

Dose distribution of OMP TPS shown less deviation compared to Monaco TPS calculated by using CC and PB algorithms. The large differences in percentage deviation between two commercial TPS (Monaco and OMP) due different system of linear accelerator used by each TPS where Monaco is from Elekta synergy system while OMP is from Primus system. For OMP TPS, the surface dose deviation of CC algorithm was 0.7% while PB algorithm was 0.6%. Therefore, the dose deviation of CC algorithm was larger compared to PB algorithm. Hence, the range of dose deviation from surface to Point A for CC algorithm (−0.1% to 0.7%) was larger compared to PB algorithm (−0.1% to 0.6%). The result of PB algorithm gave calculated value of dose closer to the measured dose compared to CC algorithm at central axis of beam in homogeneous medium [6].

### 3.3 Dose distribution with mass density correction

The result of dose deviation for a fraction and a total fraction at Point A is shown in **Error! Reference source not found.**. Noted that the negative sign represents dose calculated lower than the standard dose. CC algorithm of OMP showed higher deviation compared to Monaco. Based on **Error! Reference source not found**.(a) for axial placement of the marker, for the marker with no corrected mass density, PB algorithm (5.0 cGy) shows higher dose deviation compared to CC algorithm (3.0 cGy) for the wire. However, CC algorithm shows a higher deviation for cable (7.0 cGy) compared to PB algorithm (6.0 cGy). The same dose deviation is shown by both algorithms when plasticine (2.0 cGy) was placed on the surface. Cable gave the highest dose deviation followed by Wire and Plasticine. For OMP TPS, the dose deviation does not change when the marker was corrected with air and water mass density. When the corrected mass density applied to Monaco TPS, the dose deviation decreasing. The deviation was smallest when corrected with water mass density for cable (2.4 cGy) and plasticine (−1.3 cGy) marker. However, dose deviation for wire (1.5 cGy) was smallest when corrected by air mass density.

Based on **Error! Reference source not found**.(b) for lateral placement of marker, for the marker with no corrected mass density, CC algorithm of OMP showed higher deviation compared to Monaco and PB algorithm (6.0 cGy) showed higher deviation compared to CC algorithm (2.0 cGy) for the wire. However, CC algorithm showed a higher deviation for plasticine (7.0 cGy) and cable (7.3 cGy) compared to PB algorithm (plasticine: 6.3 cGy and cable: 6.6 cGy). Cable gave the highest dose deviation followed by Plasticine and Wire. The dose deviation for OMP TPS did not change when the marker was corrected with air and water mass density. For CC algorithm of Monaco TPS, the dose deviation decreasing after corrected with air mass density and the deviation was smallest when corrected with water mass density for wire (−0.4 cGy), cable (1.5 cGy) and plasticine (−0.5 cGy) marker.

The marker corrected with mass density during the planning was intended to reduce the dose deviation. The OMP TPS showed the same result as the marker with no corrected mass density for both algorithms. The factor related to this finding is the range of mass density in TPS (highest density: bone). For the CC algorithm of Monaco TPS, the deviation was decreased when marker corrected with air and water. For axial placement, the deviation was smallest when marker corrected with the mass density of water for cable and plasticine but, dose deviation for wire was smallest when corrected by mass density of air. For lateral placement, the deviation was smallest when corrected with the mass density of water for wire, cable and plasticine marker.

### 3.4 Dose distribution for axial and lateral placement marker

The PB algorithms gave higher dose deviation compared to CC algorithm when the wire marker was introduced on phantom surface. PB algorithm has less consideration in the change of density in dose calculation [9]. In this study, while the homogeneous medium was used, PB algorithms consider high atomic number (Z) material (radiopaque) in the calculation. For axial placement marker, Cable gave the highest dose deviation followed by Wire (mass density: 1.0973 g/cm^3^) and Plasticine. According to the calculated mass density, Plasticine with the largest mass density of 1.3956 g/cm^3^ should give the highest perturbation. However, this study showed that Cable (mass density: 1.1735 g/cm^3^) gave the highest perturbation factor. For lateral placement marker, Cable gave the highest dose deviation followed by Plasticine and Wire. In this section, the Cable also gave the highest perturbation factor. The lateral placement of marker gave smaller deviation which gave less perturbation photon beam. Therefore, the placement of marker was best at lateral orientation.

### 3.5 Dose deviation in fraction

For axial placement, the highest dose deviation per fraction using CC algorithm of Monaco TPS for the marker with no mass density correction (the standard planning used), was shown by Cable (3.4 cGy) followed by Wire (2.8 cGy) and Plasticine (2.3 cGy). The dose deviation for total fraction was Cable (54.4 cGy) followed by Wire (44.8 cGy) and Plasticine (36.8 cGy). For lateral placement, the highest dose deviation per fraction using CC algorithm of Monaco TPS for the marker with no mass density correction was shown by Cable (3.3 cGy) followed by Plasticine (2.6 cGy) and Wire (0.1 cGy). The dose deviation for total fraction was Cable (52.8 cGy) followed by Plasticine (41.6 cGy) and Wire (1.6 cGy).

## 4. Conclusion

The verification of calculated dose distribution by CC algorithm of Monaco TPS and OMP TPS and PB algorithm of OMP TPS were below the tolerance limit (2%). PB algorithm calculated the dose distribution closer to the measured dose distribution compared to the CC algorithm inhomogeneous medium. However, when the marker perturbed the photon beam, the PB algorithm gave the high deviation of the dose which conclude that the PB considered high Z material in the calculation as by CC algorithm. This study concluded that wire and plasticine were suitable to be as scar marker. This is because, even though the wire is radiopaque material and act as the standard scar marker, it has the closest mass density to water. Instead of having high mass density, plasticine as radiolucent material gave less perturbation to megavoltage photon beam. The placement of the marker in lateral orientation gave smaller perturbation to megavoltage photon beam compared to axial orientation in the overall case. Therefore, the placement of marker was best at lateral orientation.

## 5. Acknowledgement

The authors would like to extend their highest gratitude to Department of Higher Education (Malaysia), for providing a grant under RUI Grants, No:1001/CIPPT/8011001. Appreciation also goes to, the staff of the Radiotherapy Unit in Advance Medical and Dental Institute, Universiti Sains Malaysia; School of Physics, Universiti Sains Malaysia, and Siemens Medical System U.S.A

